# Heterogeneous generation of new cells in the adult echinoderm nervous system

**DOI:** 10.1101/021840

**Authors:** Vladimir S. Mashanov, Olga R. Zueva, José E. García-Arrarás

## Abstract

Adult neurogenesis, generation of new functional cells in the mature central nervous system (CNS), has been documented in a number of diverse organisms, ranging from humans to invertebrates. However, the origin and evolution of this phenomenon is still poorly understood for many of the key phylogenetic groups. Echinoderms are one such phylum, positioned as a sister group to chordates within the monophyletic clade Deuterostomia. They are well known for the ability of their adult organs, including the CNS, to completely regenerate after injury. Nothing is known, however, about production of new cells in the nervous tissue under normal physiological conditions in these animals. In this study, we show that new cells are continuously generated in the mature radial nerve cord (RNC) of the sea cucumber *Holothuria glaberrima*. Importantly, this neurogenic activity is not evenly distributed, but is significantly more extensive in the lateral regions of the RNC than along the midline. Some of the new cells generated in the apical region of the ectoneural neuroepithelium leave their place of origin and migrate basally to populate the neural parenchyma. Gene expression analysis showed that generation of new cells in the adult sea cucumber CNS is associated with transcriptional activity of genes known to be involved in regulation of various aspects of neurogenesis in other animals. Further analysis of one of those genes, the transcription factor *Myc* showed that it is expressed, in some, but not all radial glial cells, suggesting heterogeneity of this CNS progenitor cell population in echinoderms.

## 1 Introduction

In recent decades, the notion of the fixed developmental state of the adult central nervous system (CNS) has become an outdated dogma. The often life-long ability to produce new cells and incorporate them into existing circuits has now been demonstrated for a wide variety of organisms, from mammals (including humans) to invertebrates (Cayre et al., 2002; Adolf et al., 2006; Grandel et al., 2006; Fernández-Hernádndez et al., 2013; Benton et al., 2014; Urbán and Guillemot, 2014; Beltz et al., 2015; Lin and Iacovitti, 2015), yet the field of adult neurogenesis is still relatively young, with many gaps in our knowledge remaining to be filled.

For one thing, the origin and evolution of adult neurogenesis in various animal taxa remain an open question. For example, in some areas of the vertebrate brain, the ability to produce new neurons is considered a phylogenetically ancient trait, while in others, such as the dentate gyrus of mammals, it is though to be an evolutionary new adaptation (Kempermann, 2012; Urbán and Guillemot, 2014).

Another important question is whether the ability of the mature CNS to generate new cells under physiological conditions can be harnessed to repair injuries. In mammals, neural injuries result in increased proliferation in two continuously active neurogenic zones, the subventricular zone and the hippocampal dentate gyrus, and also activate quiescent neural progenitors in other brain regions (Lin and Iacovitti, 2015; Sun, 2015). This response, however, is not sufficient to fully restore the organization and function of the damaged mammalian CNS. On the other hand, there are animals who can completely repair neural injures. Further comparative studies of post-traumatic and physiological neurogenesis in those models can yield useful insights into how the limited mammalian CNS regeneration can be improved.

Echinoderms are a phylum of marine invertebrates that shares common ancestry with chordates within the monophyletic group Deuterostomia. They are well known for the unusually high regenerative capacity of their adult organs, including the CNS, and thus can provide valuable data on the evolution of neural plasticity in deuterostomes. Neurogenesis in adult echinoderms has been studied during post-traumatic regeneration (Mashanov et al., 2008; San Miguel-Ruiz et al., 2009; Mashanov et al., 2012b, 2013, 2015a,b), but little is known about generation of new CNS cells under normal physiological condition, besides the mere observation of the presence of dividing cells in the radial nerve cord (Mashanov et al., 2010, 2013). Further investigation of these endogenous progenitors present in the intact CNS will allow us to better understand the processes triggered by the injury and the nature of cell sources recruited for neural repair.

We have previously established that in both the uninjured and regenerating echinoderm CNS the majority of new cells are produced through proliferation of radial glia (Mashanov et al., 2008, 2010, 2013). Radial glial cells of echinoderms share a number of characteristics with their chordate counterparts, including some immunocytochemical properties and the typical elongated shape that allows them to span the entire height of the neuroepithelium between the apical and basal surfaces (Viehweg et al., 1998; Mashanov et al., 2006, 2009, 2010). It has remained, however, unclear whether the radial glia progenitors in all regions of the adult echinoderm CNS are equally capable of producing new cells and whether the newly born cells remained at the place of their birth or migrated to populate other regions of the CNS. In this study, we present evidence suggesting that, within the main radial nerve cords, the production of new cells is more extensive in the lateral regions than in the midline zone and that some of the cells born in the apical zone of the ectoneural epithelium might leave the place of their origin to migrate basally and populate the underlying neural parenchyma. Moreover, many of the genetic factors known to drive adult neurogenesis in vertebrates are also expressed in zones of new cell production in the adult sea cucumber CNS.

## 2 Materials and Methods

### 2.1 Animal collection, BrdU injection, and tissue sampling

Adult individuals of the sea cucumber *Holothuria glaberrima* Selenka, 1867 (Echinodermata: Holothuroidea) were collected from the rocky shore of northeastern Puerto Rico. For the duration of the experiment, the animals were kept at room temperature in indoor tanks with aerated natural sea water, which was changed weekly.

The animals were injected intracoelomically with 5-bromo-2-deoxyuridine (BrdU, Sigma), at a dose of 50 mg/kg. Injections were repeated at regular 12 h-intervals for 7 days, so that each animal received a total of 14 injections. In order to elucidate if the distribution of newly born cells varied at different time intervals, the animals were sacrificed at 4 hours (0 weeks), 1 week, 5 weeks, and 8 weeks after the last BrdU injection. Four individuals were used at each of the time points. Before dissection, the animals were anesthetized by immersion in a 0.2% chlorobutanol (Sigma) solution for 10−30 min or until they showed no response to touch.

For immunocytochemistry and in situ hybridization, pieces of the body wall containing the radial nerve cord were quickly dissected out and fixed overnight at 4°C in buffered 4% paraformaldehyde prepared in 0.01 M PBS, pH 7.4. The tissue samples were then washed in the same buffer, cryoprotected in buffered sucrose and embedded in the Tissue-Tek embedding medium (Sakura Finetek).

### 2.2 BrdU immunohistochemistry

Serial cryosections (10-*μ*m thick) were collected on gelatin-covered slides and postfixed in formalin vapors for 15 min to prevent section detachment during the subsequent staining procedure. The slides were then washed in PBS, pretreated with 0.5% Triton X-100 and incubated in 2N HCl for 30 min at 37°C to expose the BrdU epitopes in the nuclear DNA. After neutralization in 0.1M borate buffer, autofluorescence was quenched by incubation in 0.1M glycine in PBS for 1 hour. The sections were then blocked in 2% goat serum. The primary anti-BrdU antibody (1:400, GenWay, 20-783-71418) were applied overnight at 4°C. After 10 washes (10 min each) with PBS, the sections were incubated in the secondary FITC-conjugated Goat Anti-Rat antibody (1:50, GenWay, 25-787-278232) for 1 hour at room temperature. Following the final washes, the sections were mounted in a medium containing 2.5% DABCO (Sigma-Aldrich) and 10% Mowiol 4-88 (Calbiochem) dissolved in 25% glycerol buffered with 0.2M Tris-HCl (pH 8.5).

### 2.3 Cell counting

Immunostained cryosections were photographed with a Nikon Eclipse 600 microscope equipped with a SPOT RT3 camera (Diagnostic Instruments, Inc.) using a 40× objective. The acquired images were assembled into panoramic multichannel composite micrographs using the stitching plugin (Preibisch et al., 2009) in Fiji image analysis software (Schindelin et al., 2012). The cross section area of the ectoneural part of the RNC was divided into ten sampling areas as follows. The width of the RNC was divided into five areas of equal width from left to right. Each of these five areas was further subdivided into the apical zone containing dense accumulation of cell bodies and the basal zone, which included the neural parenchyma (Figs. 1, 2). All clearly BrdU-labeled cells (strongly and moderately stained) were counted on every third cross-section, five sections per animal, using the Cell Counter plugin in Fiji. The total number of BrdU^+^-cells was divided by the total area of the corresponding sampling region to calculate the of BrdU^+^-cell density (Additional File 1).

**Figure 1:**
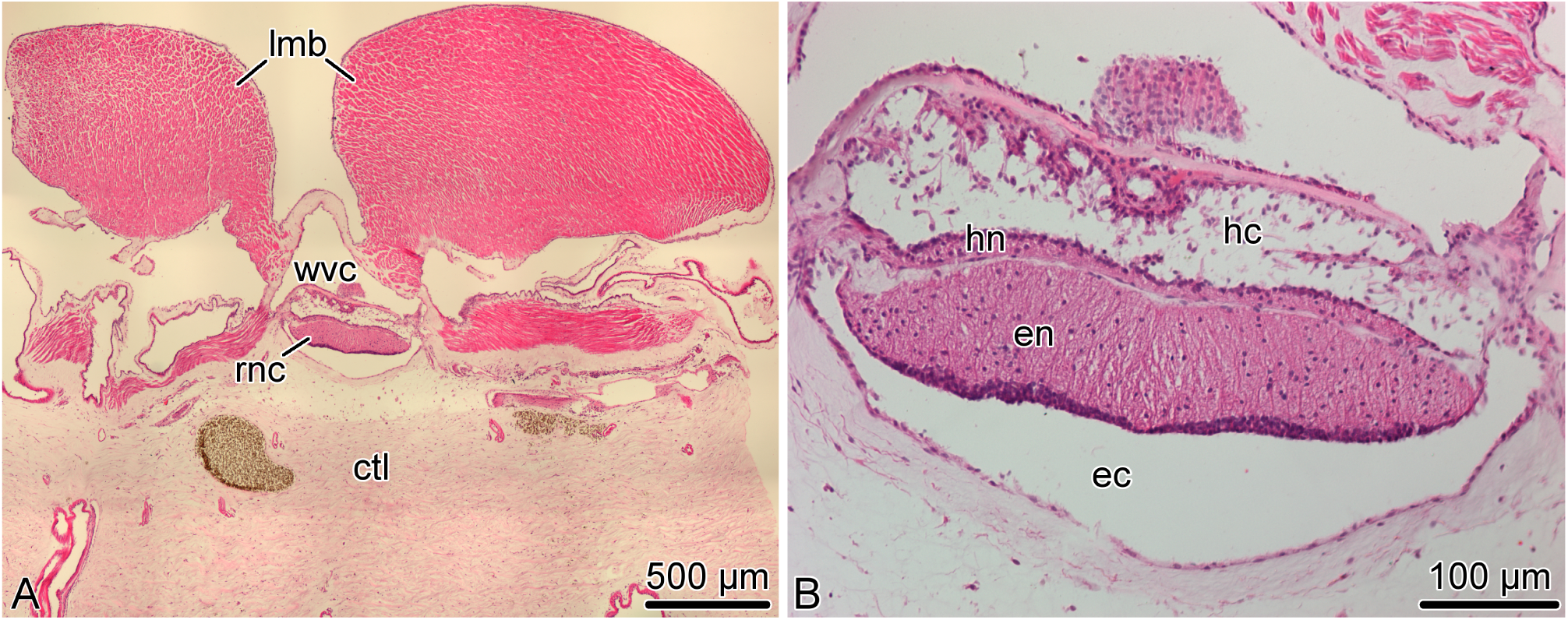
Organization of the radial nerve cord (RNC) in the sea cucumber *H. glaberrima*. (A) Low magnification overview of a cross section of the body wall showing the position of the radial nerve cord *(rnc)* relative to other anatomical structures, such as the longitudinal muscle band *(lmb)*, radial canal of the water-vascular system *(wvc)*, and the connective tissue layer of the body wall *(ctl)*. (B) Higher magnification view of the radial nerve cord. Note two parallel bands of nervous tissue, a thicker ectoneural neuroepithelium *(en)* and a thinner hyponeural epithelium *(hn)* separated by a thin connective tissue partition. The apical surface of the ectoneural and the hyponeural canals form the bottom of the epineural *(ec)* and hyponeural *(hc)* canals, respectively. Paraffin sections; hematoxylin and eosin staining.

**Figure 2:**
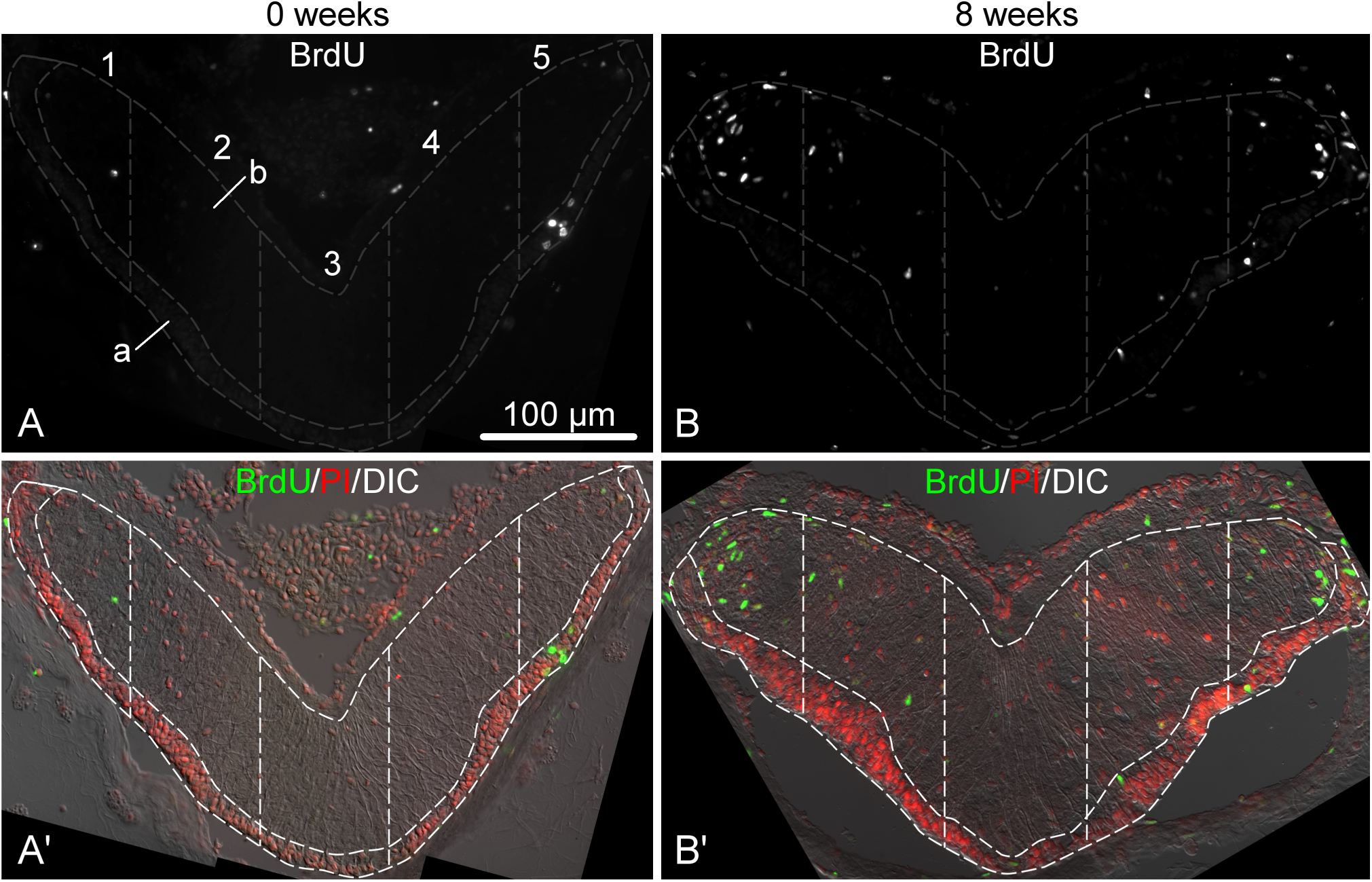
Representative micrographs showing distribution of BrdU-positive cells in the ectoneural epithelium of the RNC sampled immediately after the last BrdU injection (A and A’) and after 8 weeks (B and B’). A and B show labeling with the anti-BrdU antibody in a single channel. A’ and B’ are corresponding multichannel composite micrographs, which, besides BrdU labeling, also include nuclei labeled with propidium iodide (PI) and differential interference contrast (DIC) image of the RNC. Sampling areas for cell counting are marked with dashed lines. Along the left-right axis, the RNC was divided into five columns of equal width. Each column was then subdivided into the apical (a) and basal (b) parts corresponding the the zones of dense cell body accumulation and neural parenchyma, respectively.

### 2.4 Statistical analysis

The data were found to be non-normally distributed (right skewed). Therefore, to analyze them, we used a generalized linear modeling approach with a quasipoisson error distribution instead of classic parametric tests. All computations were performed in R (v3.1.2) (R Core Team, 2015). The statistical significance of the main effects and interactions between them were determined by F-test in analysis of deviance. Multiple pairwise comparisons between individual means were performed with the post-hoc Tukey test using the 

~~~
glht
~~~

 function of the 

~~~
multicomp
~~~

 package. The input file containing the raw data and the sample R code that can be used to reproduce the calculations can be found in Additional File 1 and Additional File 2, respectively. The full output of the statistical analysis is available in Additional File 3.

### 2.5 Gene sequences and phylogenetic analysis

Sea cucumber orthologs of genes known to be involved in neurogenesis in other animals (Table 1) were identified by TBLASTN search against the *H. glaberrima* reference transcriptome database (http://dx.doi.org/10.6070/H4PN93J1) (Mashanov et al., 2014). Sequences of *H. glaberrima Myc, Klf1/2/4, Oct1/2/11,* and *SoxB1* were retrieved from the GenBank (accession numbers KM281936−KM281939), as they have been previously characterized elsewhere (Mashanov et al., 2015a).

**Table 1.**
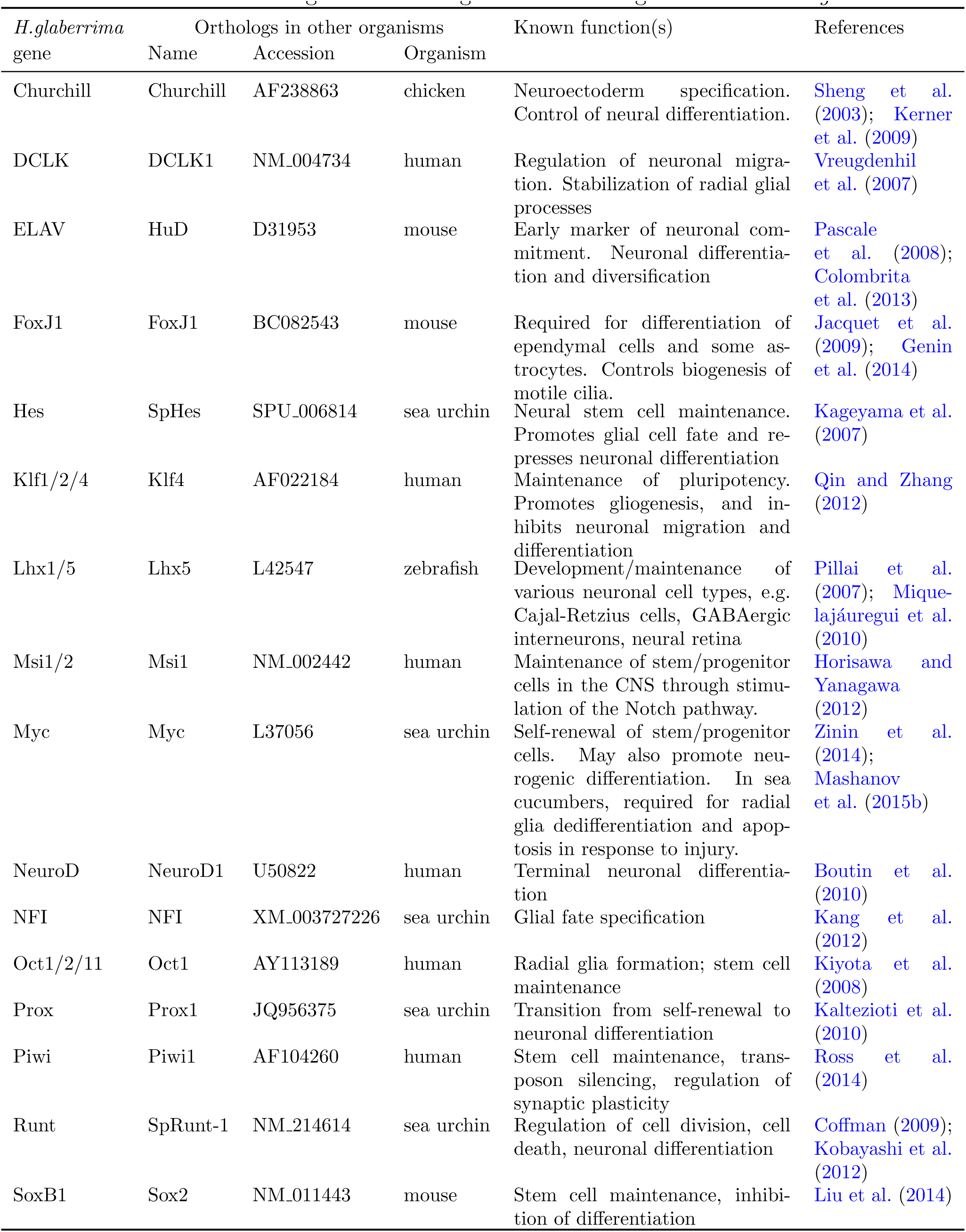
Homologs of neurogenesis-related genes in *H. glaberrima*

The identity of the sea cucumber sequences was confirmed by the analysis of the conserved domain composition through the Pfam (http://pfam.xfam.org/) database search and by analysis of phylogenetic relationships with similar proteins from other organisms (Additional File 4). These reference sequences were retrieved by BLAST search from the UniProt, EchinoBase (http://www.echinobase.org), and NCBI’s nr databases. Multiple sequence alignments of DNA coding regions were performed with ClustalW or MUSCLE and used as input to generate phylogenetic trees with MEGA (v 6.0) (Tamura et al., 2013) using the neighbor-joining method and the bootstrap test with 2,000 replicates.

### 2.6 In situ hybridization

Antisense DIG-labeled riboprobes were transcribed from PCR-generated DNA templates using Roche DIG-labeling mix (see Additional File 5 for PCR primer sequences). Tissue samples were fixed and processed for cryosectioning as described above. Frozen sections were collected onto coverslips pretreated with 3-aminopropyltriethoxysilane (APES, Sigma). Chromogenic hybridization reactions were performed in 24-well tissue culture plates as described previously (Mashanov et al., 2012a). Briefly, the sections were pretreated with proteinase K and then acetylated. Riboprobes were diluted to a final concentration of about 400 ng/ml in a hybridization buffer containing 50% formamide, 5× SSC (standard saline citrate), 0.1% Tween20, and 40 *μ*g/ml denatured salmon sperm DNA. Hybridization was performed overnight at 45°C. Following extensive stringency washes, the sections were blocked in Blocking Solution (Roche) and then incubated overnight at 4°C in alkaline-phosphatase-conjugated anti-DIG antibodies (1:2,000, Roche). After washing off the excess antibody, the color was developed in NBT/BCIP solution in the dark. After that, the sections were briefly postfixed in 4% paraformaldehyde and mounted in buffered gelatin/glycerol.

Control DIG-labeled riboprobes were synthesized from the pSPT18-Neo (DIG RNA Labeling kit, Roche) and GFP (green fluorescent protein) DNA templates. These control probes did not yield any detectable signal under the same hybridization conditions as above.

For double fluorescent labeling with the *Myc* riboprobe and the ERG1 antibody, in situ hybridization was performed first. It was carried out as above, except that the NBT/BCIP mixture was replaced with Vector Red alkaline phosphatase substrate (Vector Labs). The ERG1 monoclonal antibody, which specifically labels radial glial cells in the echinoderm nervous system (Mashanov et al., 2010), was applied overnight at 4°C. After washing off the unbound primary antibody, the sections were incubated in the Cy3-conjugated goat anti-mouse secondary antibody (1:2,000, Jackson ImmunoResearch Laboratories, Inc) for 1 hour at room temperature.

## 3 Results

### 3.1 Organization of the radial nerve cords in echinoderms

The most prominent components of the nervous system in adult echinoderms are five radial nerve cords, which are joined by a circumpharyngeal nerve ring at the oral side of the body to form an anatomically continuous CNS (Hyman, 1955). In sea cucumbers, each of the radial nerve cords (RNCs) is composed of two closely apposed layers of neural tissue called the ectoneural and hyponeural bands (Fig. 1) (Mashanov et al., 2006; Hoekstra et al., 2012). Both components of the RNC have a neuroepithelial organization. The framework of the neuroepitelium is made up of radial glial cells, whose apical cell bodies are joined together by intercellular junctions, while long basal processes span the height of the underlying neural parenchyma and attach to the basal lamina (Mashanov et al., 2006, 2010). Neuronal perikarya are scattered throughout the height of the neuroepithelium. Some of them are interspersed between the apical cell bodies of radial glia, while others are immersed into the underlying parenchyma being surrounded by zones of neuropil. The two neuroepithelia form a bottom of an epineural and hyponeural canal, respectively (Fig. 1B). The roof of the canals is a simple epithelium composed of flattened glial cells. The ectoneural neuroepithelium is the predominant component of the radial nerve cord in sea cucumbers. It is much taller than the hyponeural neuroepithelium, contains more cells, and will be the primary focus of the present study.

### 3.2 Newly born cells accumulate at different rates in different regions of the adult sea cucumber radial nerve cord

Newly born cells within the adult echinoderm CNS were identified as cells retaining bromodeoxyuridine (BrdU) immunoreactivity. Our preliminary experiments (not shown) demonstrated that a single BrdU injection followed by a 4 hour-long chase period labels only very rare cells (0.45 ± 0.10%, mean ± standard error) in the RNC, suggesting that only a limited number of cells undergoes pre-mitotic DNA synthesis at any given moment. In order to increase the number of labeled cells (to 2.60 ± 0.55%, mean ± standard error), we performed saturating BrdU injections twice a day for seven days and then monitored the distribution of labeled cells within the RNC at various time points after the last injection. Figure 2 shows representative micrographs of immunostained sections of the RNC sampled immediately after the last BrdU injection and 8 weeks post-injection. For the purpose of cell counting, the cross-sectional area of the ectoneural part of the RNC was divided into five sampling zones of equal width along the left-right axis (see the Materials and Methods section). Within each of those five regions, the neuroepithelium was further subdivided along the natural landmark separating a narrower apical zone, which contained densely packed cell bodies forming a discrete layer, and a wider basal zone, which contained the neuropil area with more loosely arranged cell bodies.

Our statistical analysis (Additional File 3) involving generalized linear modeling approach with time, apical-basal position, and left-right position as factors, showed that the density of BrdU^+^-cells varied significantly (F-test, *P* = 1.6 × 10^-11^) along the left-right axis of the ectoneural layer of the radial nerve cord, with the lateral areas (area 1 and area 5) containing significantly more labeled cells per *μ*m^2^ of cross-sectioned area than the midline area (area 3) (Fig 3). For example, eight weeks after the last BrdU injection, the mean density of BrdU-labeled cells in the lateral regions is about 2.4−3 times greater than in the midline region. There was no interaction between the mediolateral position and the other two factors, suggesting that the observed pattern of distribution of BrdU-labeled cells along the left-right axis does not change with time or with position along the apical-basal axis.

**Figure 3:**
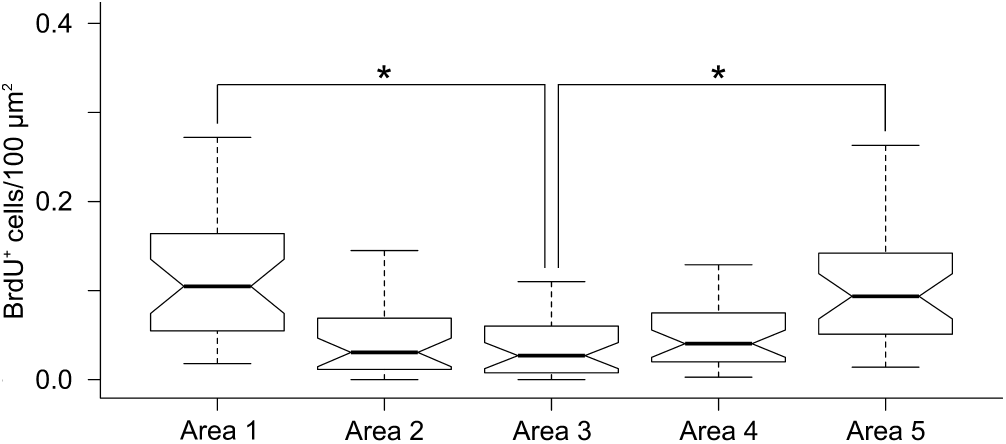
Lateral regions of the RNC have significantly higher density of BrdU^+^ cells than the mid-line regions. Notched box and whisker plots. Boxes show the interquartile range (the values between the 25% and 75% percentiles), the line within the box is the median of the data, and the whiskers represent adjacent values within the 1.5 × interquartile range outside the box. * *P <* 0.05

However, there was a significant (*P* = 1.8 × 10^-6^) interaction between the post-injection time span and the apical−basal position. We, therefore, studied the simple main effect of length of time after the last BrdU injection on the density of labeled cells separately in the apical and basal region of the RNC. There were no significant changes in the density of labeled cells in the apical region of the ectoneural neuroepithelium (F-test, *P* = 0.24; Fig. 4A) sampled at different post-injection time points. Surprisingly, however, the basal region showed significant accumulation of BrdU^+^-cells with time (F-test, *P* = 5.86 × 10^-6^; Fig. 4B). For example, the mean density of labeled cells in the neural parenchyma is ∼6.3 times greater after 8 weeks post-injection than immediately after the last injection.

**Figure 4:**
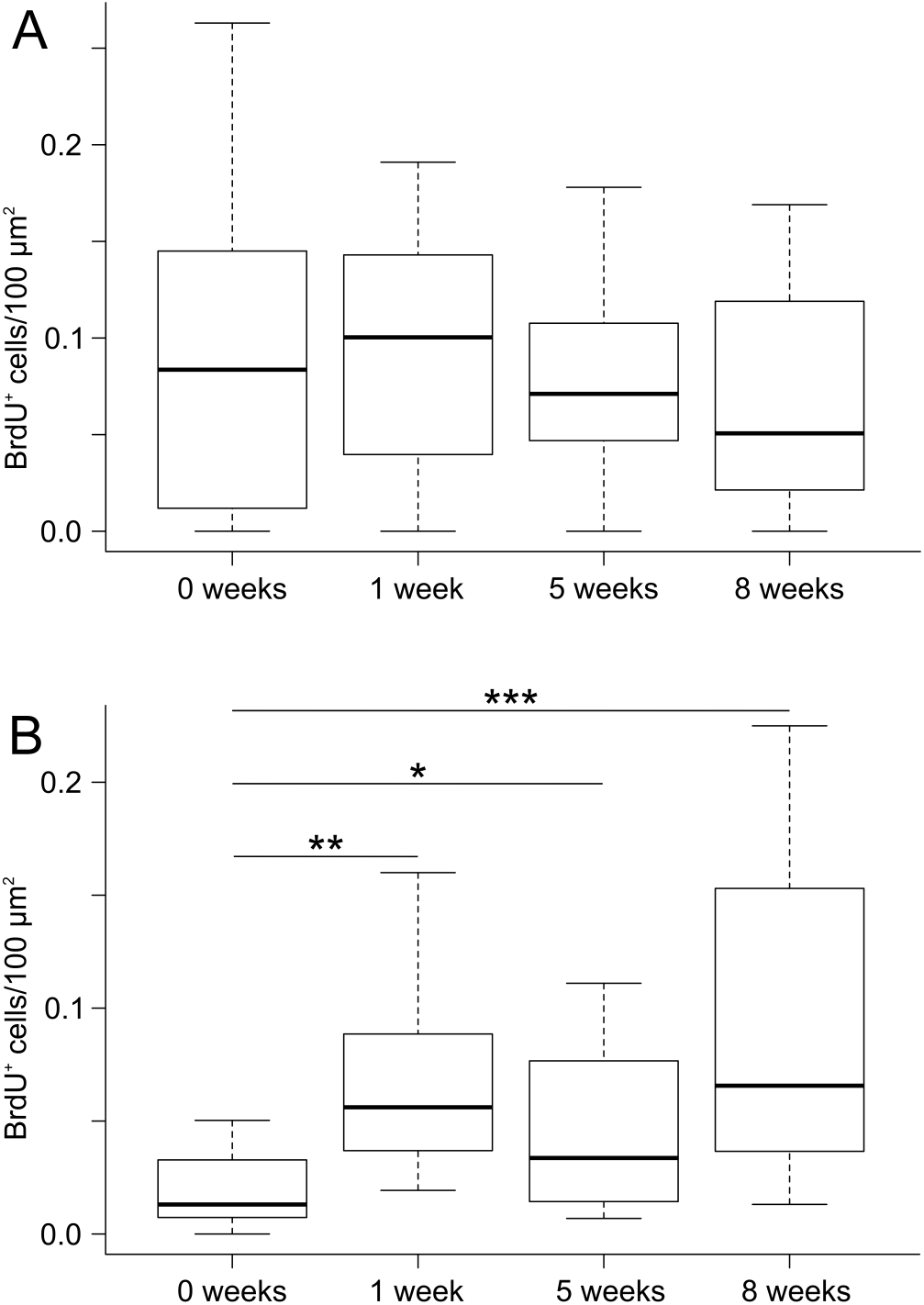
Box and whisker plot showing density of BrdU+ cells in the apical (A) and basal(B) regions of the ectoneural neuroepithelium of the RNC as a function of the length of time after the last BrdU injection. * *P <* 0.05; ** *P <* 0.01; *** *P <* 0.001

Comparison of the apical and basal zones of the ectoneural epithelium shows a significantly higher (*∼*4.7-fold, *P* = 2.9 × 10^5^ density of the labeled cells in the apical zone immediately after the last BrdU injection (Fig. 5A), but this difference between the two regions decreases with time and becomes negligible (*P* = 0.18) after 8 weeks (Fig. 5B).

**Figure 5:**
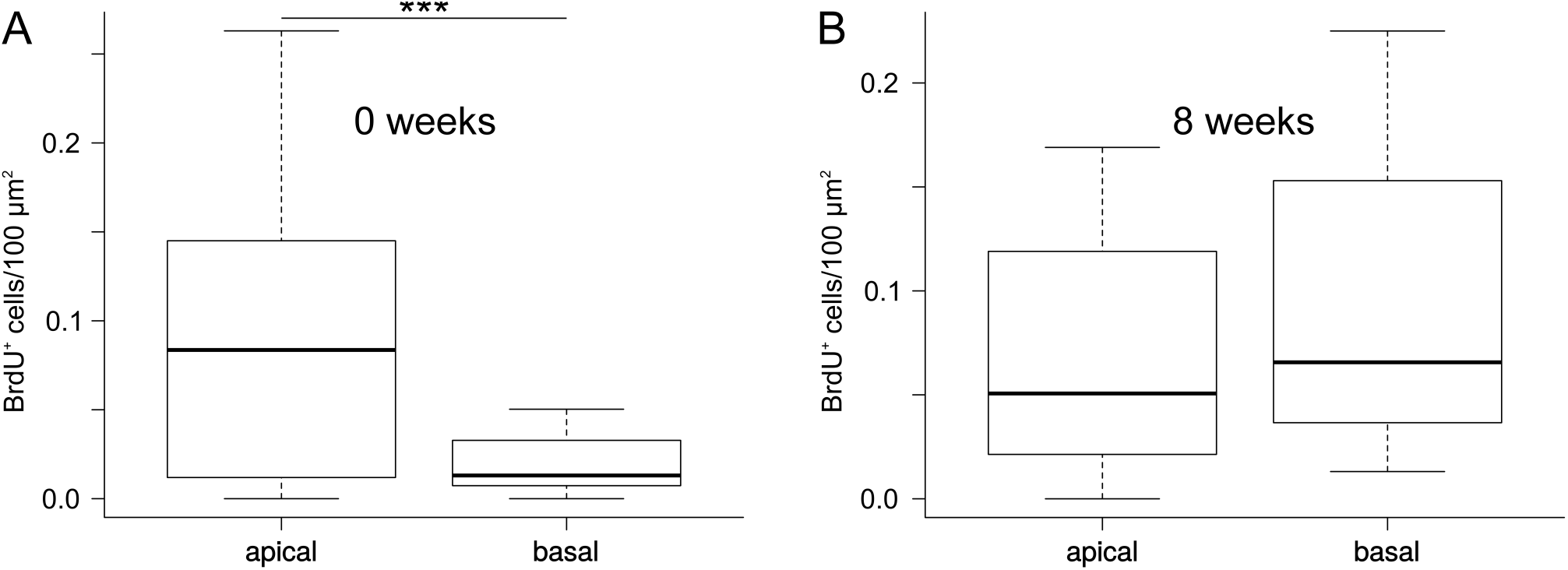
Box and whisker plots showing comparison of density of BrdU-labeled cells between the apical and basal regions of the RNC immediately after the last BrdU injection (A) and after 8 weeks (B).

Taken together, the cell counting data suggest that the production of new cells is more extensive in the lateral regions of the RNC than in the midline zone. Of note, after the last BrdU injection the density of newly produced cells remains constant in the apical region of the neuroepithelium, while steadily increasing in the basal region. We suggest that some of the BrdU-labeled cells in the apical zone undergo further cell divisions during the chase period, thus producing daughter cells that still contain enough BrdU to be identified as BrdU-positive. Some of these apical cells eventually leave the site of their origin and migrate into the basal neural parenchyma, thus keeping the abundance of the apical BrdU^+^-cells at the constant level while leading to accumulation of labeled cells in the basal region in the absence of BrdU.

### 3.3 Generation of new cells in the adult radial nerve cord is associated with expression of known neurogenic markers

The molecular mechanisms controlling cell dynamics in the adult echinoderm CNS remain completely unknown. In order to establish whether or not the production of new cells in the adult echinoderm CNS is associated with expression of known neurogenic genes, we searched the sea cucumber transcriptome database (Mashanov et al., 2014) with TBLASTN for homologs of proteins with documented roles in neural development and/or adult neurogenesis. In total, 16 genes were retrieved (Table 1), whose orthologs are involved in various aspects of neurogenesis, including maintenance of neural stem/progenitor cells (e.g., *Hes, Klf1/2/4, Msi1/2, Myc, Oct1/2/11, Piwi,* and *SoxB1*), neural differentiation (e.g., *Churchill, ELAV, Lhx1/5, Myc, NeuroD, Prox,* and *Runt*), and glial fate specification (*FoxJ1, Klf1/2/4,* and *NFI*). With the exception of the previously studied (Mashanov et al., 2015a) four genes, *Klf1/2/4, Myc, Oct1/2/11*, and *SoxB1*, the *H. glaberrima* database sequences returned by the BLAST search are contigs derived from automated de novo assembly of next generation sequencing reads. We, therefore, validated them by re-sequencing with Sanger technology and deposited the verified sequences in GenBank under accession numbers KT183348 − KT183359. The orthology between these newly characterized sea cucumber genes and the corresponding genes in other organisms was confirmed by phylogenetic analysis (Figs. 6−8). We then employed in situ hybridization in order to investigate spatial expression of the above genes at the tissue level in the adult sea cucumber RNC (Figs. 9−11). All 16 genes are strongly and extensively expressed in the apical region of the ectoneural neuroepithelium. However, this expression domain is discontinuous, as a narrow band of midline cells shows no positive staining with any of the riboprobes used in this study. In addition to being expressed in the apical region of the neuroepithelium, ten of the above genes, including *Hes, Klf1/2/4, Myc, SoxB1, Piwi, NFI, DCLK, Prox, Runt*, and *Lhx1/5* are also expressed in scattered cells in the neural parenchyma. The labeled parenchymal cells are most abundant in the lateral regions of the RNC and are rare in the midline region (see e.g., Fig. 9C, D, G, G” and Fig. 11A, A”, F, F”).

**Figure 6:**
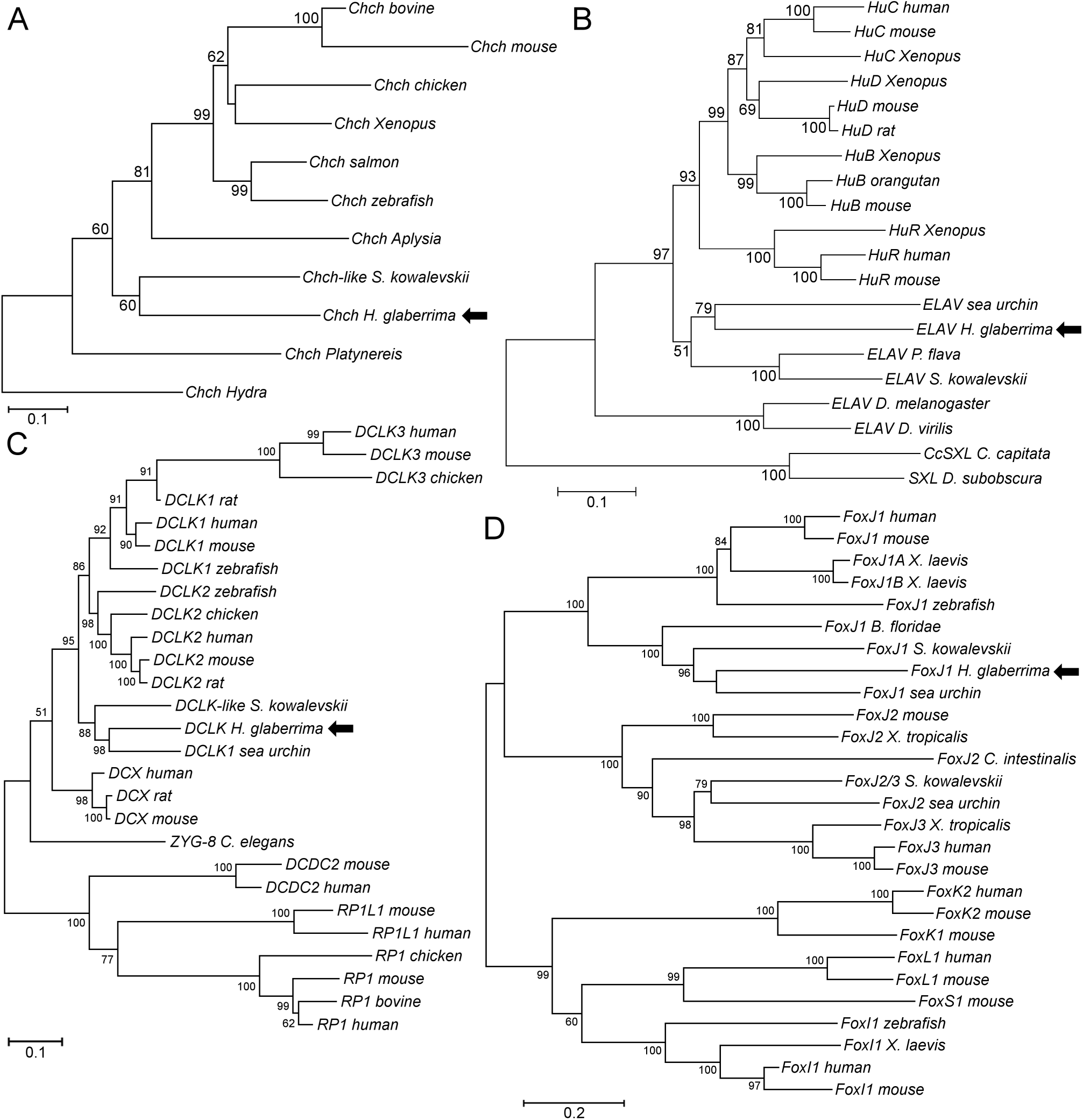
Neighbor-joining trees showing phylogenetic relationships of *H. glaberrima Churchill* (A), *ELAV* (B), *DCLK* (C), and *FoxJ1* (D) *(arrows)* with homologous genes from other organisms. Bootsrap values higher than 50% (2,000 replicates) are shown next to the branches.

**Figure 7:**
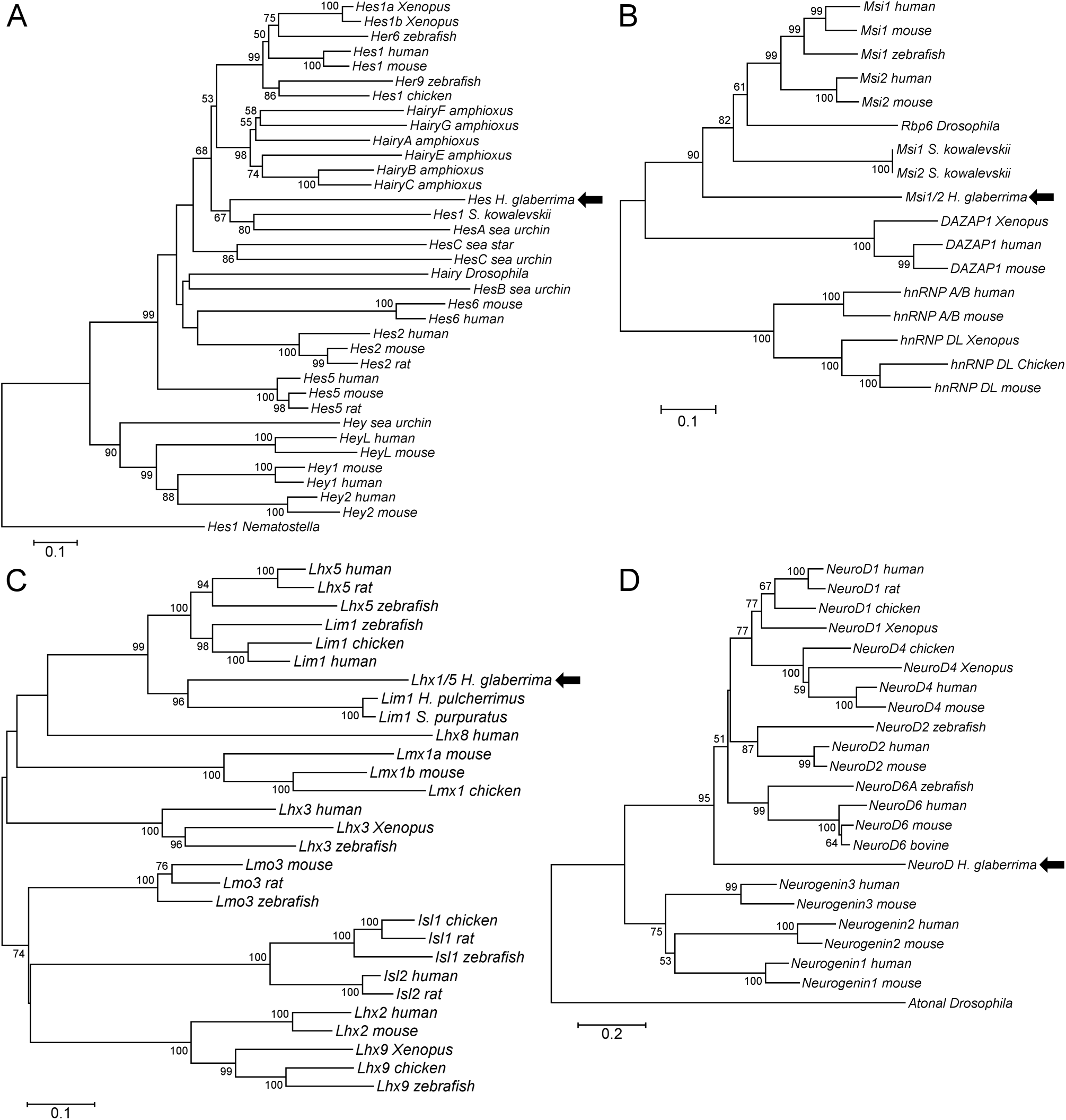
Neighbor-joining trees showing phylogenetic relationships of *H. glaberrima Hes* (A), *Msi1/2* (B), *Lhx1/5* (C), and *NeuroD* (D) *(arrows)* with homologous genes from other organisms. Bootsrap values higher than 50% (2,000 replicates) are shown next to the branches.

**Figure 8:**
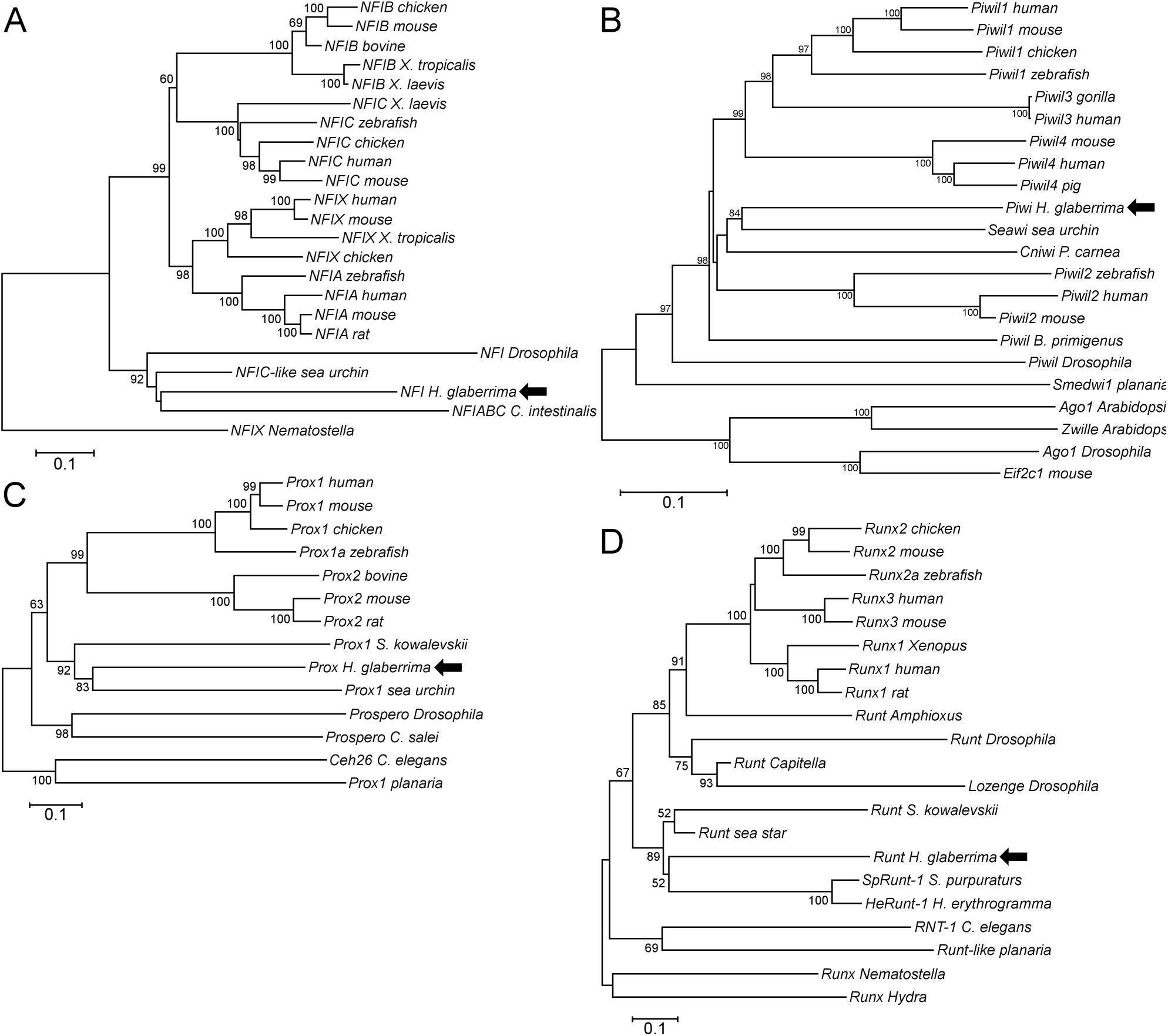
Neighbor-joining trees showing phylogenetic relationships of *H. glaberrima NFI* (A), *Piwi* (B), *Prox* (C), and *Runt* (D) *(arrows)* with homologous genes from other organisms. Bootsrap values higher than 50% (2,000 replicates) are shown next to the branches.

**Figure 9:**
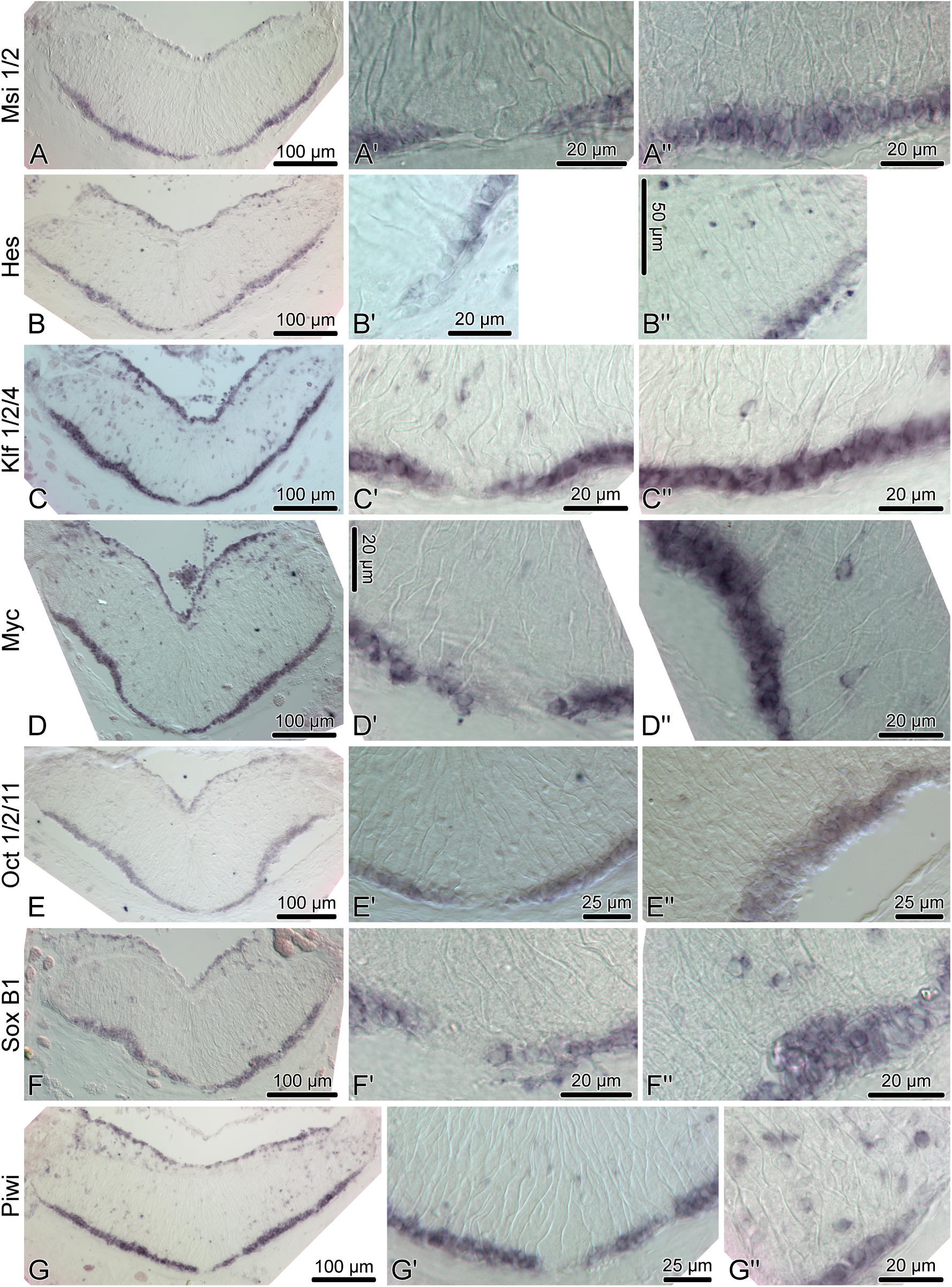
Expression of homologs of neural stem/progenitor cell maintenance genes in the adult radial nerve cord (RNC) of *H. glaberrima*. The left column shows reference low magnification micrographs of the entire cross section profile of the RNC. The middle column is a detailed view of the midline region of the ectoneural neuroepithelium. Micrographs in the right column are higher magnification of the lateral region of the ectoneural neuroepithelium.

Among the genes that show robust expression in the apical zone, but little or no expression the neural parenchyma, are *ELAV* (Fig. 11B−B”) and *NeuroD* (Fig. 11C−C”), two factors involved in neuronal differentiation (Pascale et al., 2008; Boutin et al., 2010), suggesting that some processes involved in neuronal maturation can take place only at the apical surface of the neuroepithelium, but not in the deeper parenchyma.

**Figure 11:**
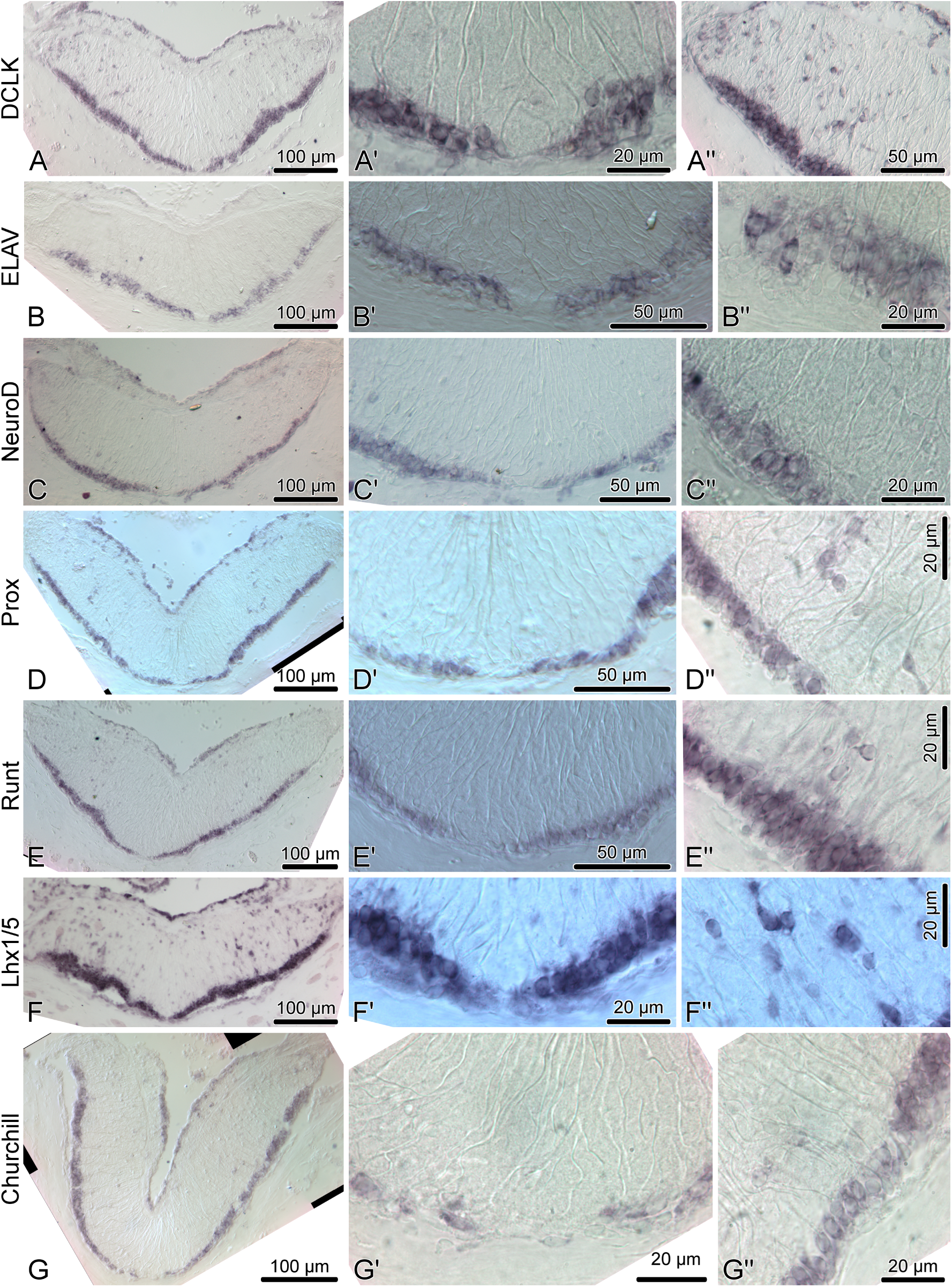
Expression of proneural genes in the adult radial nerve cord (RNC) of *H. glaberrima*. The left column shows reference low magnification micrographs of the entire cross section profile of the RNC. The middle column shows a detailed view of the midline region of the ectoneural neuroepithelium. Micrographs in the right column are higher magnification of the lateral region of the ectoneural neuroepithelium.

Another transcription factor with exclusive expression in the apical zone of the ectoneural neuroepithelium is *FoxJ1* (Fig. 10A−A”). This expression pattern correlates well with the known role of this gene in ciliogenesis (Jacquet et al., 2009; Genin et al., 2014). The apical cell bodies of echinoderm radial gial cells have been previously shown to bear a cilium protruding into the lumen of the epineural canal (Mashanov et al., 2006).

**Figure 10:**
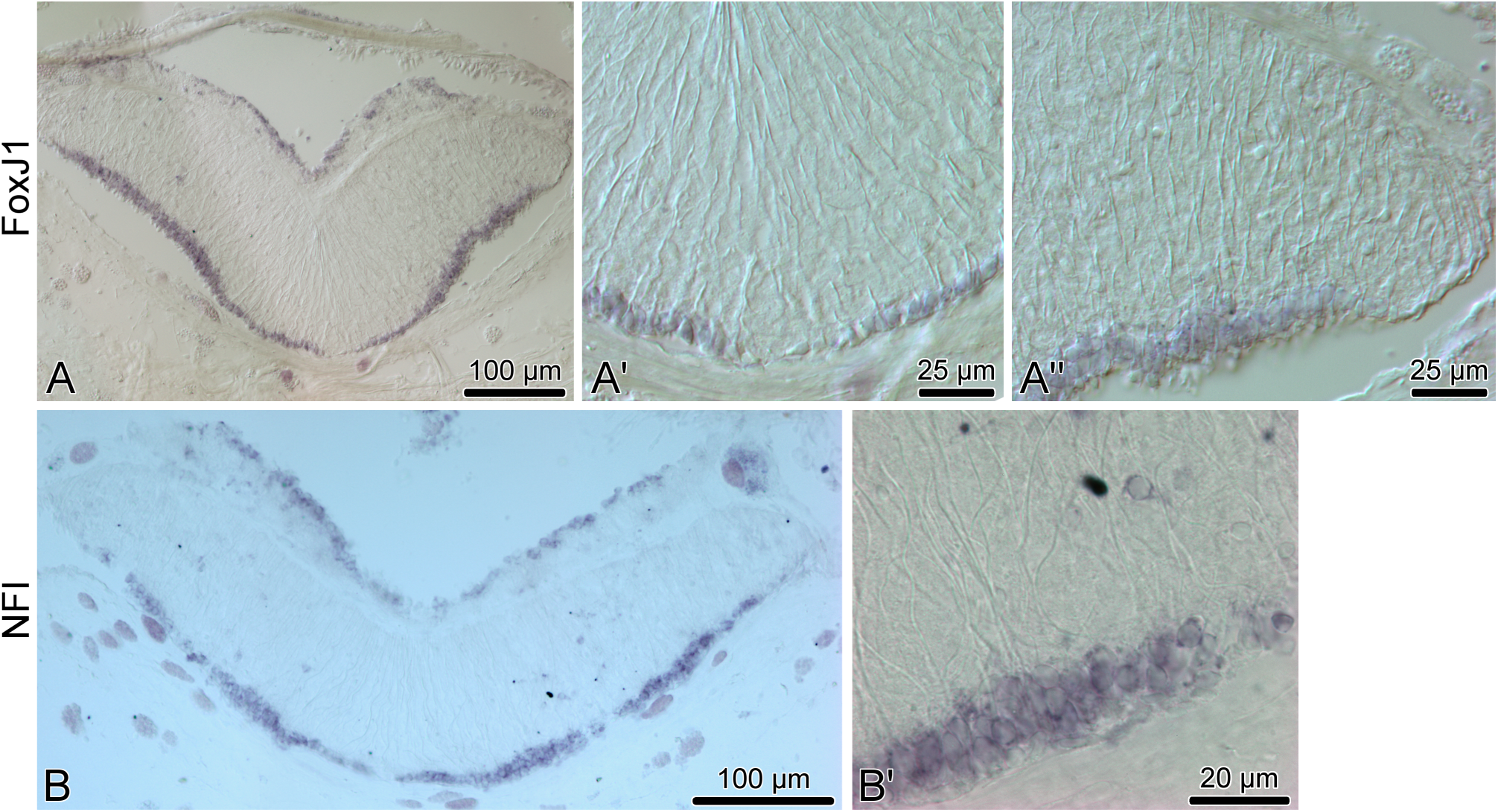
Expression of proglial genes, *FoxJ1* (A−A”) and *NFI)* (B and B’), in the radial nerve cord (RNC) of *H. glaberrima*. (A) and (B) show low magnification overview micrographs of the entire cross section profile of the RNC. (A’) is a detailed view of the midline region of the ectoneural neuroepithelium. (A”) and (B’) show higher magnification views of the lateral region of the ectoneural neuroepithelium.

### 3.4 Echinoderm radial glial cells are heterogeneous

Our earlier studies (Mashanov et al., 2010, 2013) identified radial glial cells as a major progenitor cell population in the adult echinoderm CNS, as they were shown to account for the majority of cell divisions in the nervous tissue both under physiological conditions and after injury. Most importantly, at least some of the progeny of these proliferating cells can differentiate into neurons.

Cells bodies of most of the radial glial cells lie in the apical region of the neuroepithelium (Mashanov et al., 2006, 2010), i.e., within the domain of most robust expression of all sixteen neurogenesis-related genes described in the previous section. However, it has never been directly demonstrated that the radial glial cells of adult echinoderms express any of these neurogenesis-related genes at the single cell level under physiological conditions. Moreover, it is not known whether or not the echinoderm radial glial cells, which are all morphologically alike, are homogeneous in terms of their expression profile.

In order to provide at least a partial answer, we simultaneously labeled the radial nerve cord with a specific glial marker, the ERG1 monoclonal antibody (Mashanov et al., 2010), and with a *Myc* in situ hybridization riboprobe. The choice of *Myc* was determined by phylogenetically conserved role of Myc proteins in activation of neural progenitors, which was demonstrated in animals as diverse as rat and *Drosophila* (Hasegawa et al., 2005; Fernández-Hernández et al., 2013). In the regenerating echinoderm radial nerve cord, *Myc* was also recently shown to be required for glial activation in response to injury (Mashanov et al., 2015b). In this study, the double labeling showed that many of the radial glial cells indeed expressed *Myc*, but these positively labeled cells were interspersed with glial cells, which showed no *Myc* expression (Fig. 12). These results suggest that the echinoderm radial glial cells, despite being all morphologically alike, differ in expression of at least some of the key transcription factors. It remains to be determined whether or not this heterogeneity in gene expression results in heterogeneity in potency and the ability to proliferate.

**Figure 12:**
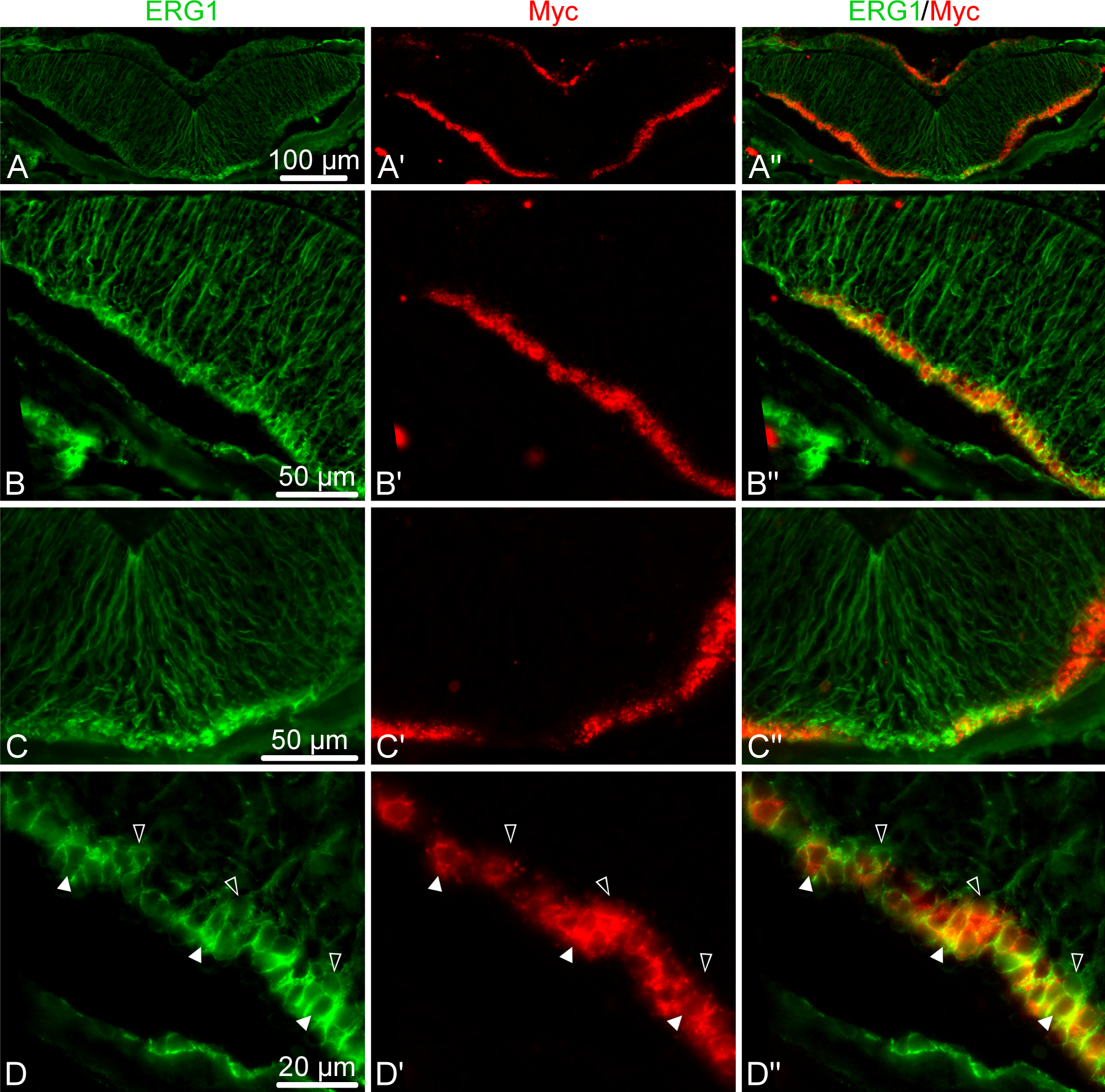
Double fluorescent labeling with the ERG1 antibody, a maker of echinoderm radial glial cells (Mashanov et al., 2010) (green, left column), and an in situ hybridization probe for *Myc* (red, middle column). The right column shows overlay composite images with both markers. (A−A”) Low-magnification of a cross section through the radial nerve cord. (B−B”) Detailed view of the lateral region of the ectoneural neuroepithelium. (C−C”) Detailed view of the midline region of the ectoneural neuroepithelium. (D−D”) High-magnification view of the apical region of the ectoneural epithelium showing colocalization of the *Myc* in situ signal with ERG1 labeling in cell bodies of some of the radial glial cells *(white arrows)*, whereas other glial cells do not express *Myc* at all *(open arrows)*

## 4 Discussion

The ability to generate new cells in the adult CNS is widespread in the animal kingdom and has been documented in both vertebrates and invertebrates. In mammals, adult neurogenesis is mostly restricted to two distinct brain regions, the subventricular zone of the anterior ventricles and the subgranular zone of the hippocampal dentate gyrus (Urbán and Guillemot, 2014; Lin and Iacovitti, 2015). In the CNS of non-mammalian vertebrates, such as fish, proliferation sites are more abundant, but are still confined to certain discrete loci (Adolf et al., 2006; Grandel et al., 2006). To our knowledge, nothing has been known so far about the pattern of new cell generation in the adult CNS of more basal deuterosomes. The data presented in this paper suggest that the RNCs of echinoderms are heterogeneous too in their ability to produce new cells, mostly clearly along the mediolateral axis. The regions where newly generated cells are most abundant are organized into longitudinal domains along the lateral sides of the nerve cords. The physiological significance of restricting production of new cells to these lateral regions remains to be elucidated.

Another interesting parallel with neurogenesis in vertebrates is that at least some of the new cells generated in the sea cucumber RNC migrate away from the place of their birth. Along the apical-basal axis of the ectoneural neuroepithelium, one can distinguish the apical domain, where the density of BrdU-positive cells remains constant throughout long chase periods, and the basal domain with the steadily increasing abundance of BrdU-positive cells. The most parsimonious explanation of the observed data is that the labeled cells may keep dividing in both zones, but some of the cells born in the apical region of the neuroepithelium migrate into the underlying basal zone, thus keeping the labeled cell number constant in the former and leading to its rise in the latter.

Although many of the transcription factors known to control neurogenesis in other animals have been identified in the sea urchin genome and were studied in association with the developing larval nervous system (Burke et al., 2006; Poustka et al., 2007), little is known about genetic mechanism controlling adult neurogenesis in echinoderms. In this study, we show that production of new cells in the sea cucumber CNS is associated with expression of sixteen genes, whose homologs are known to control various aspects of vertebrate neurogenesis, including neurogenic stem cell maintenance, neuronal specification and glial differentiation. Strong steady expression of a combination of pluripotency factor homologs, including *SoxB1, Oct1/2/11, Klf1/2/4*, and *Myc*, as well as of other stem cell-associated genes, such as *Piwi*, in the apical zone of the ectoneural epithelium suggests that this region of the adult sea cucumber CNS contains cells that constantly retain progenitor properties.

The above genes, however, are not necessarily homogeneously expressed in all cells throughout the apical region of the neuroepithelium, not even in cells belonging to the same cell type. We have previously demonstrated that radial glial cells, the only major glial cell type in echinoderms, accounts for almost all cell divisions in both the normal and regenerating adult CNS (Mashanov et al., 2010, 2013) and thus acts as the key progenitor cell population. Morphologically, all radial glial cells in echinoderms have the same organization. They are all tall slender cells stretched between the apical and basal surfaces of the neuroepithelium. Most of the radial glial cells are unipolar, with an apically positioned nucleus and a long basal process (Mashanov et al., 2006, 2010). Nevertheless, in this study we showed that in spite of morphological homogeneity, the echinoderm radial glial cells are distinctly heterogeneous in their expression of the pluripotency factor *Myc*. Heterogeneity of CNS stem/progenitor cells in terms of their gene expression profile, activation/quiescence status, commitment to the generation of specific progenitors, etc. is a common property of adult neurogenesis in various vertebrate species (Urbán and Guillemot, 2014; Lin and Iacovitti, 2015; Than-Trong and Bally-Cuif, 2015). It remains, however, to be elucidated if the heterogeneity in expression of *Myc*, and possibly other transcription factors, in the sea cucumber radial glia results in differences in the capacity to generate new cells in the adult CNS.

Among other genes, whose expression is associated with generation of new cells in the adult sea cucumber CNS, are also regulators and effectors of the Notch signaling pathway. Notch is expressed in radial glia and neuronal progenitors in zebrafish, and in neural stem cells in mammals. Acting through *Hes* genes, Notch signaling regulates maintenance of neural progenitors and represses their premature differentiation (Kageyama et al., 2007; Urbán and Guillemot, 2014; Than-Trong and Bally-Cuif, 2015). In the adult sea cucumber *H. glaberrima*, *Notch* transcripts are present in both the uninjured and regenerating RNC (Mashanov et al., 2014). Here, we show that *Hes*, a direct target of Notch signaling, is prominently expressed in the apical zone of the ectoneural epithelium and in some scattered cells in the neural parenchyma, suggesting involvement of active Notch signaling in sea cucumber neurogenesis. *Msi* and *Elav*, the positive and negative modulators of Notch signaling, respectively (Glazer et al., 2008; Horisawa and Yanagawa, 2012), share the same expression domain with *Hes*, suggesting a precise control of the balance between cell proliferation and differentiation.

## 5 Conclusions

Our study demonstrates the following

- New cells are being constantly produced in the adult echinoderm radial nerve cord.
- Newborn cells are significantly more abundant in the lateral region of the radial nerve cord, than along the midline
- Some of the cells produced in the apical region of the ectoneural neuroepithelium leave their place of birth to migrate into the underlying neural parenchyma.
- Generation of new cells in the adult sea cucumber CNS is associated with expression of genes whose orthologs are implicated in control of various aspects of neurogenesis in other animals
- In spite of stereotypical morphology, at the single cell level the radial glial cells, the major progenitor cell population in the echinoderm CNS, are heterogenous in terms of gene expression. We show that radial glial cells expressing the transcription factor *Myc* are interspersed with glial cells that do not express this gene.

## 6 Acknowledgments

The study was supported by grants from the NIH (1R03NS065275-01) and the NSF (IOS0842870, IOS-1252679), as well as by the University of Puerto Rico.

## 8 Additional Files

### Additional File 1

Comma-separated file containing the raw measurements of density of BrdU-labeled cells (number of BrdU^+^-cells per *μ*m^2^) in the ectoneural epithelium of *H. glaberrima*

### Additional File 2

R code used to perform statistical computations.

### Additional File 3

Analysis of deviance table showing the effects of the length of post-injection time, position along the apical-basal axis and left-right axis on the density of BrdU-labeled cells in the ectoneural neuroepithelium of the radial nerve cord of the sea cucumber *H. glaberrima*

### Additional File 4

Reference numbers of reference sequences used to generate phylogenetic trees

### Additional File 5

Sequences of PCR primers used to generate templates for riboprobe synthesis

